# Arf GTPases Define BST-2-Independent Pathways for HIV-1 Assembly and Release

**DOI:** 10.1101/2025.07.28.667227

**Authors:** Adam Smith, Dominique Dotson, Jessica Sutton, Hua Xie, Xinhong Dong

## Abstract

ADP-ribosylation factor (Arf) proteins are small GTPases that regulate intracellular membrane trafficking and actin cytoskeleton remodeling through cycles of GTP binding and hydrolysis. Arf1, a central regulator of Golgi and endosomal trafficking, and Arf6, which controls plasma membranes and endosomal dynamics, have both been implicated in late stages of the HIV-1 life cycle. However, the precise mechanisms by which these GTPases support HIV-1 assembly and release remain incompletely understood. Here, we provide direct evidence that Arf1 and Arf6 are required for efficient trafficking of the HIV-1 Gag polyprotein, assembly, and virion release. Perturbation of Arf1 function with either a GTP-locked (Q71L) or GDP-locked (T31N) mutant significantly reduced virus release, impaired Gag association with membrane compartments, and blocked Gag targeting to the plasma membrane. Manipulation of Arf1 activity via the GTPase-activating protein AGAP1 further demonstrated that dynamic cycling of Arf1 between GTP- and GDP-bound states is essential for productive Gag trafficking. Similarly, expression of a constitutively active Arf6 mutant (Q67L) misdirected Gag to intracellular membranes and markedly decreased virion production. Importantly, disruption of Arf1 or Arf6 function did not affect the total expression, surface levels, or intracellular distribution of the host restriction factor BST-2. Together, these findings establish that Arf1- and Arf6-mediated trafficking pathways are critical host determinants of HIV-1 assembly and release, functioning independently of BST-2 antagonism.

**IMPORTANCE:** The small GTPases Arf1 and Arf6 control fundamental processes in membrane trafficking and cytoskeletal dynamics, yet their roles in HIV-1 replication are not well defined. We show that both proteins are required for efficient trafficking of HIV-1 Gag polyprotein to the plasma membrane and for subsequent virus release. Disrupting either GTPase reroutes Gag to intracellular membranes and reduces virion production, independently of the antiviral host factor BST-2. These results identify Arf1- and Arf6-dependent trafficking as critical host pathways for HIV-1 assembly and egress, expanding our understanding of the cellular machinery hijacked by retroviruses to support infections.

## INTRODUCTION

The Gag protein is the major structural component of HIV-1 and other retroviruses, orchestrating virion assembly, release, and maturation (1–3). Remarkably, Gag alone is sufficient to drive the formation and release of virus-like particles (VLPs) from the cell surface in the absence of other viral components. HIV-1 Gag is synthesized as a polyprotein precursor comprising four major structural domains: the N-terminal matrix (MA), capsid (CA), nucleocapsid (NC), and C-terminal p6 domains (1,4). Each domain plays distinct roles during late stages of the viral life cycle (5,6). The MA domain, through a N-terminal myristylation signal and a highly basic region (HBR), directs membrane targeting and binding (7,8). The CA domain governs Gag-Gag interactions required for multimerization (9,10), while the NC domain binds and packages the viral RNA genome and also contributes to Gag multimerization during assembly (11). The p6 domain recruits the host ESCRT (endosomal sorting complexes required for transport) machinery through interaction with Tsg101 and ALIX, thereby enabling membrane fission and virion release (12–14).

While the intrinsic biochemical properties of Gag are sufficient for particle formation in heterologous expression systems, increasing evidence highlights the importance of host cell trafficking pathways and cytoskeletal networks in regulating Gag localization and virion egress (15–18). Small GTPases of the ADP-ribosylation factor (Arf) family are key regulators of membrane traffic and vesicle biogenesis, particularly in the context of Golgi and endosomal transport (19–21). Arf proteins cycle between GDP-bound (inactive) and GTP-bound (active) states to coordinate cargo sorting, vesicle formation, and organelle identity (22,23). Among these, Arf1 Arf6 have emerged as regulators of retroviral egress, although the precise mechanisms remain incompletely defined (24,25). Arf1 predominantly localized to the Golgi and regulates vesicle trafficking via adaptor proteins such as the GGAs (Golgi-localized, γ-ear-containing, Arf-binding proteins), while Arf6 functions at the plasma membrane and endosomal compartments, controlling actin dynamics, endocytosis, and recycling (26,27). Both Arf1 and Arf6 have been implicated in the release of retroviruses such as HIV-1, murine leukemia virus (MLV), and equine infectious anemia virus (ELAV), raising the possibly that Arf-mediated trafficking influences viral assembly or access to key egress factors (24).

One such egress factor is BST-2 (bone marrow stromal antigen 2; also known as tetherin or CD317), an interferon-inducible type II transmembrane protein that potently inhibits the release of HIV-1 and other enveloped viruses. BST-2 functions by physically tethering nascent virions to the plasma membrane or to each other, preventing their dissemination from infected cells (28–30). Structurally, BST-2 consists of an N-terminal cytoplasmic tail, a single-pass transmembrane domain, a coiled-coil extracellular domain, and a C-terminal glycosylphosphatidylinositol (GPI) anchor (31) . This unique topology allows it to insert one membrane anchor into the plasma membrane and the other into budding virions, thereby forming a physical bridge that retains virions at the cell surface (28,32,33). In response, HIV-1 encodes the accessory protein Vpu, which antagonizes BST-2 by promoting its degradation through both proteasomal and lysosomal pathways, sequestering it in intracellular compartments, and displacing it from viral budding sites (34–36). The Vpu-BST-2 axis represents a key example of host-pathogen antagonism and serves as a model for understanding how viruses counteract intrinsic immunity.

While both Arf proteins and BST-2 have been independently linked to HIV-1 egress, it remains unclear whether Arf-mediated trafficking regulates the localization, stability, or antiviral function of BST-2. Moreover, whether the impact of Arf proteins on HIV-1 release is dependent on BST-2 antagonism or reflects parallel trafficking mechanisms required for Gag transport remains to be determined.

In this study, we investigated the role of Arf1 and Arf6 in modulating HIV-1 particle production, Gag subcellular localization, and BST-2 trafficking. Using biochemical, virological, and imaging approaches, we demonstrate that both Arf1 and Arf6 are required for efficient trafficking of HIV-1 Gag to the plasma membrane and for productive virus release. Disruption of either GTPase alters Gag trafficking, leading to intracellular accumulation and substantial reduction in VLP production. Importantly, these effects are independent of BST-2 expression or localization, suggesting that Arf1 and Arf6 promote HIV-1 assembly and egress through BST-2-independent pathways. These findings identify Arf GTPases as critical host factors co-opted by HIV-1 to support late stages of the viral life cycle, likely by directing Gag trafficking through distinct membrane compartments en route to the plasma membrane.

## RESULTS

### Arf1 regulates HIV-1 particle release and Gag subcellular localization

To define the role of the small GTPase Arf1 in HIV-1 particle production, HEK 293T cells were co-transfected with plasmids encoding HIV-1 Gag-Pol and either an empty vector (pcDNA3.1), wild-type (WT) HA-tagged Arf1, a constitutively active GTP-locked mutant (HA-Arf1/Q71L), or a dominant-negative GDP-locked mutant (HA-Arf1/T31N). At 48 h post-transfection, cell lysates and pelleted virus-like particles (VLPs) from clarified supernatants were analyzed by immunoblotting with HIV-IgG to detect Gag proteins. Overexpression of WT Arf1 did not significantly alter Gag expression, processing or VLP release compared to the vector control. In contrast, expression of either Arf1/Q71L or Arf1/T31N moderately but reproducibly reduced VLP release (Fig 1A), indicating that dynamic cycling between GTP-and GDP-bound states is required for efficient HIV-1 particle production. To quantify the extent of this effect, VLP release efficiency was calculated as the ratio of VLP-associated p24 to total p24 (cell-associated plus VLP-associated) and normalized to the vector control (Gag-Pol + pcDNA3.1, lane 1).

**Fig. 1.**
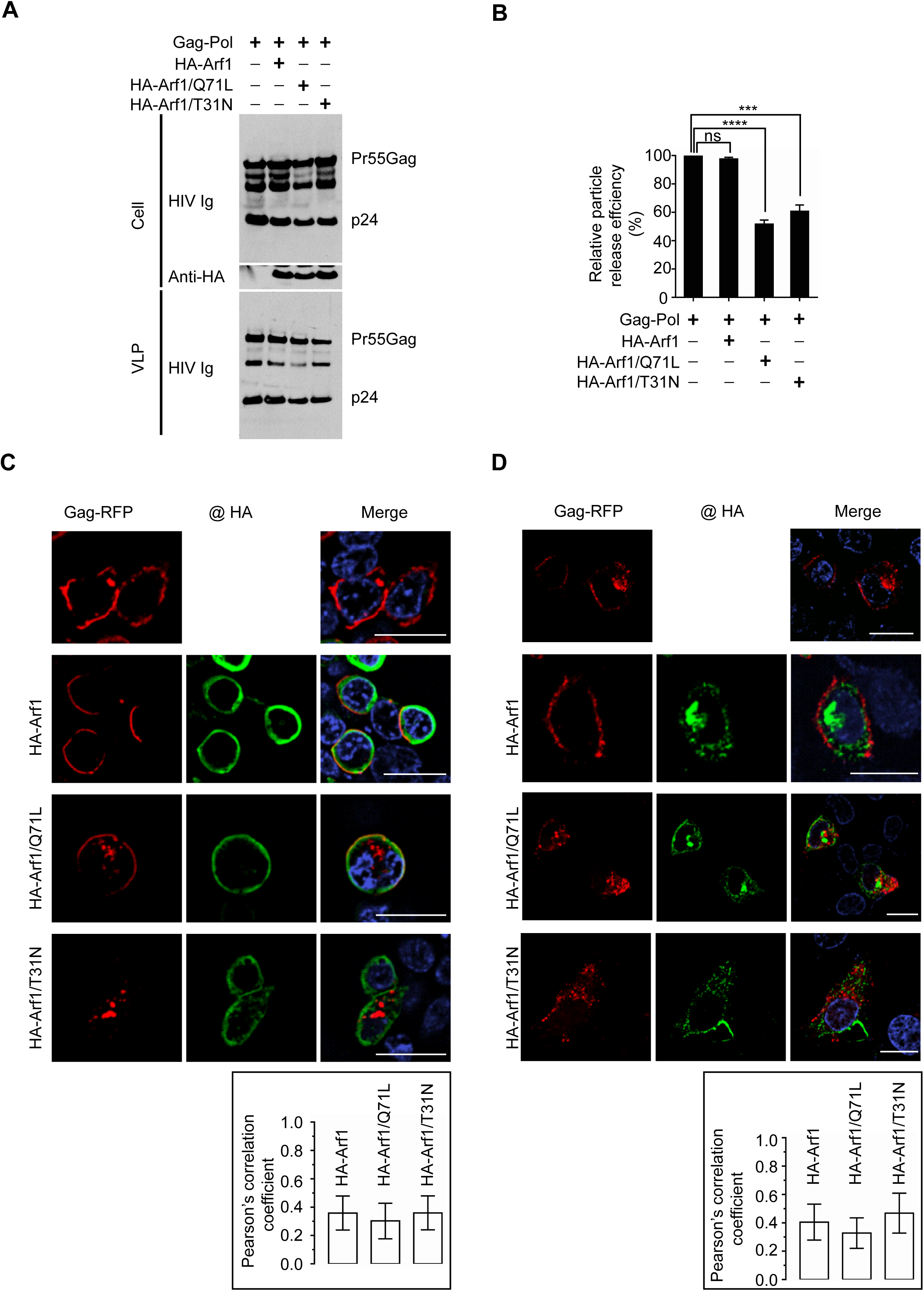
Arf1 regulates HIV-1 particle production and subcellular localization of Gag. **A,** HEK 293T cells were co-transfected with HIV-1 Gag-Pol and either empty vector (pcDNA3.1, lane 1), WT HA-Arf1 (lane 2), HA-Arf1/Q71L (lane 3), or HA-Arf1/T31N (lane 4) at a 1:1 plasmid ratio. Cell lysates were analyzed by immunoblotting using HIV-IgG and anti-HA antibodies, and VLPs pelleted from the supernatant were probed with HIV-IgG. **B,** Quantification of relative particle release efficiency from (A). Release efficiency was calculated as the amount of p24 in the supernatant divided by total p24 (supernatant plus cell-associated), and normalized to the pcDNA3.1 control. Data represent mean ± SD from six independent experiments (n=6). ns, not significant; ***, *p* < 0.001; ****, *p* < 0.0001. **C-D,** Confocal microscopy of HEK 293T (C) or HeLa (D) cells expressing Gag-RFP alone (top row), Gag-RFP with HA-Arf1 (second row), Gag-RFP with HA-Arf1/Q71L (third row), or Gag-RFP with HA-Arf1/T31N (bottom row). Gag-RFP is shown in red (left panels), HA-tagged proteins in green (middle panels), and merged images in yellow (right panels). Scale bars, 20 μm. Pearson’s correlation coefficient (R) were used to quantify colocalization. Graphs show mean ± SD from 20-30 cells per condition, based on ≥ 3 independent experiments.

Expression of either Arf1/Q71L or Arf1/T31N reduced VLP release efficiency by approximately 50%, whereas WT Arf1 had no significant effect (Fig. 1B). These results indicate that both impaired GTP hydrolysis (Q71L) and defective GTP loading (T31N) compromise Arf1 function during HIV-1 assembly and/or release.

To further examine how Arf1 activity affects Gag trafficking, we performed confocal microscopy in HEK 293T cells co-expressing Gag-RFP and HA-tagged Arf1 constructs. In cells expressing Gag-RFP alone, the protein localized predominantly to the plasma membrane, consistent with its role in viral assembly. Co-expression with WT HA-Arf1 did not alter this pattern; Gag-RFP remained membrane-associated, and minimal colocalization was observed between Gag and Arf1, which localized to submembranous regions. In contrast, expression of either HA-Arf1/Q71L or HA-Arf1/T31N disrupted Gag-RFP trafficking, leading to reduced plasma membrane accumulation and increased localization to perinuclear compartments (Fig. 1C). Both Arf1 mutants retained a submembranous distribution similar to WT Arf1. Quantitative colocalization analysis using Pearson’s correlation coefficients revealed low overlap between Gag-RFP and HA-Arf1 in all conditions: 0.36 ± 0.12 for WT, 0.30 ± 0.13 for Q71L, and 0.36 ± 0.12 for T31N. To further resolve these trafficking defects, similar experiments were conducted in HeLa cells, which display more defined intracellular membrane architecture. In control cells, Gag-RFP localized to both the plasma membrane and internal compartments. However, expression of either Arf1/Q71L or Arf1/T31N markedly reduced Gag-RFP at the plasma membrane and redirected it to perinuclear regions (Fig.1D), recapitulating the phenotype observed in HEK 293T cells. Pearson’s correlation coefficients in HeLa cells remained low to moderate: 0.40 ± 0.12 for WT, 0.33 ± 0.11 for Q71L, and 0.46 ± 0.14 for T31N. To confirm these observations, we performed parallel experiments using untagged Gag instead of Gag-RFP. In these experiments, Gag was detected using anti-p17 (MA) antibodies. Gag alone predominantly localized to the plasma membrane at 24 h post-transfection. However, co-expression of either HA-Arf1/Q71L or HA-Arf1/T31N resulted in Gag accumulation at internal membrane and a corresponding reduction in plasma membrane localization (Fig. S1). Together, these findings demonstrate that perturbation of Arf1 GTPase cycling impairs Gag trafficking and reduces HIV-1 particle release, highlighting a functional requirement for dynamic Arf1 activity in the late stages of viral assembly and budding.

### Arf1 regulates the membrane association of HIV-1 Gag

To determine whether Arf1 influences the membrane association of HIV-1 Gag, we performed membrane flotation assays using discontinuous iodixanol gradients. HEK 293T cells were co-transfected with plasmids encoding HIV-1 Gag and either empty vector (pcDNA3.1), WT HA-Arf1, HA-Arf1/Q71L, or HA-Arf1/T31N. At 48 h post-transfection, cells were harvested, lysed in hypotonic buffer, and post-nuclear supernatants were subjected to ultracentrifugation on 10-50% iodixanol step gradients. Twelve fractions were collected from top (low density, enriched in membrane-associated material) to bottom (high density, enriched in cytosolic or membrane-free components) and analyzed by immunoblotting using p24 to detect Gag and anti-HA to detect Arf1 variants. In cells expressing Gag alone, Gag was distributed across both membrane-associated fractions (fractions 2-5) and denser, membrane-free fractions (fractions 8-12) (Fig. 2A). Co-expression with WT HA-Arf1 did not significantly alter this distribution pattern (Fig. 2B). In contrast, cells expressing either HA-Arf1/Q71L or HA-Arf1/T31N resulted in a pronounced shift of Gag away from membrane-associated fractions and into denser fractions (Fig. 2C, 2D), consistent with impaired membrane targeting. Immunoblotting of total lysates confirmed comparable expression levels of Gag and Arf1 constructs across conditions (Fig. 1A), indicating that altered subcellular distribution was not due to differences in protein abundance.

**Fig. 2.**
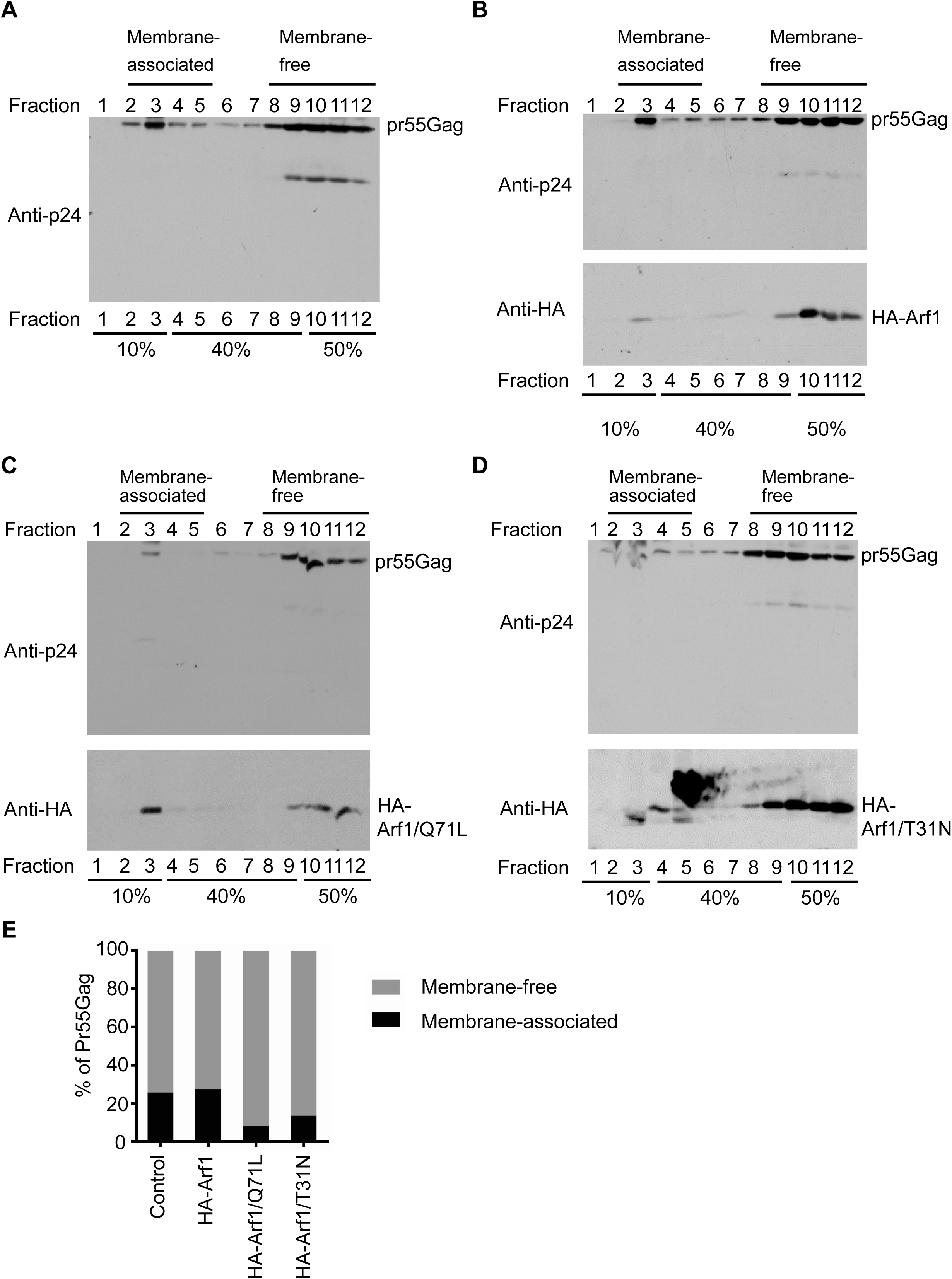
Arf1 regulates the membrane association of HIV-1 Gag. **A-D,** HEK 293T cells were co-transfected with HIV-1 Gag and either (A) empty vector (pcDNA3.1), (B) WT HA-Arf1, (C) HA-Arf1/Q71L, or (D) HA-Arf1/T31N. At 48 h post-transfection, cells were lysed and subjected to membrane flotation centrifugation. Twelve equal-volume fractions were collected from the top of the gradient and analyzed by immunoblotting with anti-p24 and anti-HA antibodies. Factions 2-5 correspond to membrane-associated proteins, while fractions 8-12 represent cytosolic (membrane-free) proteins. **E,** Quantification of membrane-associated versus membrane-free Gag distribution across conditions. Bar graphs represent the percentage of Gag in membrane-associated (fractions 2-5) and membrane-free (fractions 8-12) fractions. Data are representative of three independent experiments.

Quantitative analysis of Gag distribution revealed that the proportion of Gag in fractions 2-5 was reduced by ∼50% in cells expressing either Arf1 mutant compared to vector or WT Arf1 controls (Fig. 2E). This reduction correlated with the decrease in VLP release efficiency observed in parallel experiments (Fig. 1B), suggesting that Arf1 promotes HIV-1 egress by facilitating Gag localization to appropriate membrane compartments. The fact that both the GTP-locked (Q71L) and GDP-locked (T31N) mutants disrupted Gag membrane association supports the model that dynamic cycling between GFP- and GDP-bound states is essential for Arf1 function in this context.

### AGAP1 modulates HIV-1 particle release and Gag trafficking

Because Arf1 function is regulated by GTPase-activating proteins (GAPs), we next investigated the role of AGAP1, an Arf1-specific GAP, in HIV-1 particle production (37,38). HEK 293T cells were co-transfected with HIV-1 Gag-Pol and either WT FLAG-tagged AGAP1 or a GAP-deficient mutant (FLAG-AGAP1/R599K) at plasmid ratios ranging from 1:1 to 1:2 relative to Gag-Pol. Total DNA input was equalized with the addition of empty vector (pcDNA3.1) as needed. Immunoblot analysis of cell lysates and pelleted VLPs showed that WT AGAP1 expression reduced VLP release in a dose-dependent manner (Fig. 3A), consistent with the hypothesis that excessive Arf1 inactivation interferes with HIV-1 assembly or budding. In contrast, expression of the catalytically inactive AGAP1/R599K mutant had minimal effect on VLP production. Quantification of p24 antigen levels in cells and supernatants confirmed a significant reduction in VLP release efficiency in cells expressing WT AGAP1, but not in those expressing AGAP1/R599K (Fig. 3B). These results support a model in which AGAP1 regulates HIV-1 egress by promoting proper Arf1 GTPase cycling, and that disruption of this regulatory mechanism impairs particle release.

**Fig. 3.**
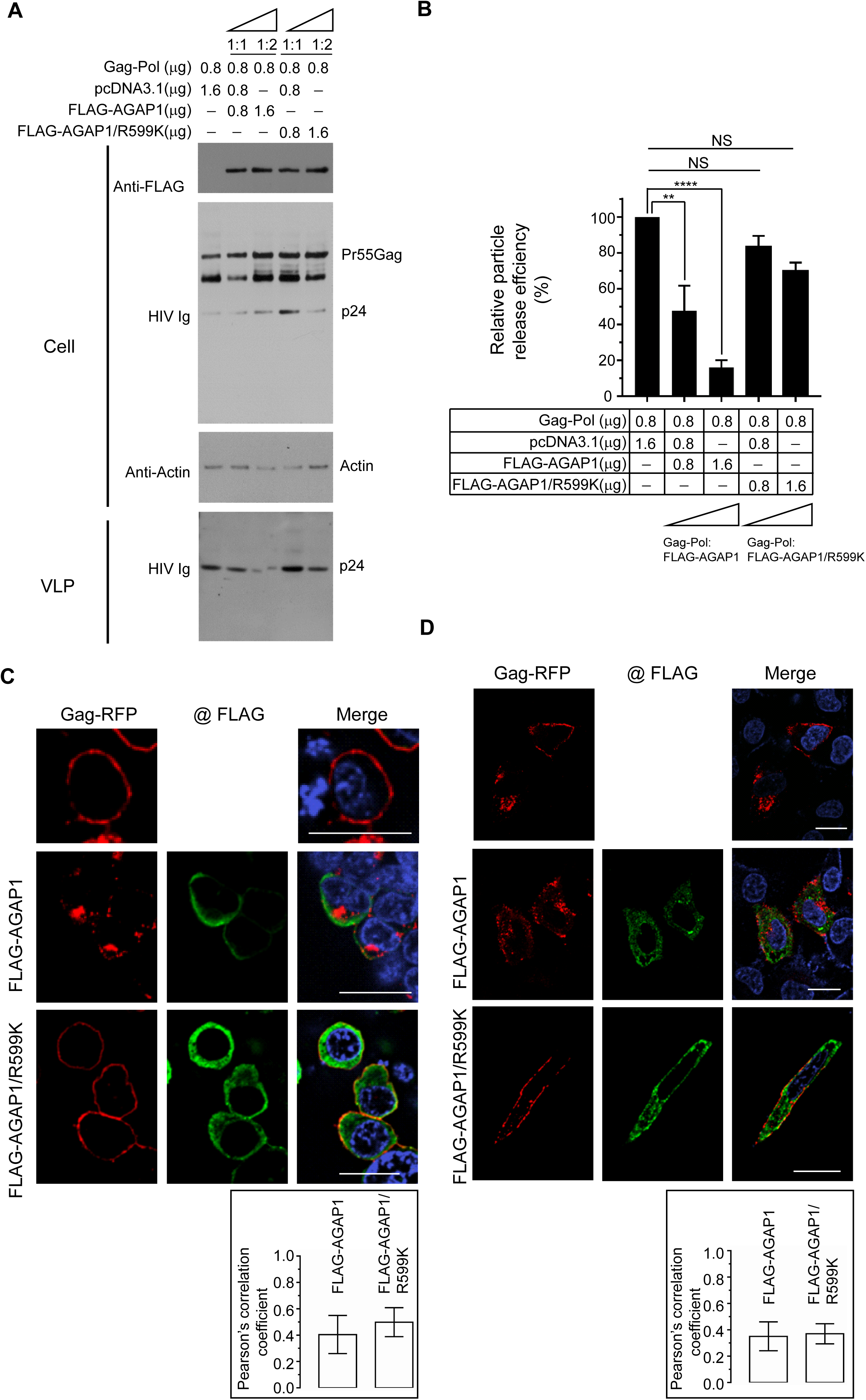
AGAP1 modulates HIV-1 particle release. **A,** HEK 293T cells were co-transfected with HIV-1 Gag-Pol and either WT FLAG-AGAP1(lanes 2-3), or FLAG-AGAP1/R599K (lanes 4-5) at increasing plasmid ratios (from 1:0 to 1:2 relative to Gag-Pol). Empty vector (pcDNA3.1) was used to equalize total DNA input across samples. Cell lysates were analyzed by immunoblotting using anti-FLAG, HIV-IgG, and anti-actin antibodies. Pelleted VLPs were probed with HIV-IgG. **B,** Quantification of relative particle release efficiency from (A). Release efficiency was calculated as the ratio of p24 in the VLP fraction to total p24 (VLP plus cell-associated), normalized to the control. Data represent mean ± SD (n=3). ns, not significant; **, *p* < 0.01; ****, *p* < 0.0001. **C-D,** Confocal microscopy of HEK 293T (C) and HeLa (D) cells expressing Gag-RFP alone (top row), Gag-RFP with FLAG-AGAP1 (second row), or Gag-RFP with FLAG-AGAP1/R599K (bottom row). Gag-RFP is shown in red (left panels); FLAG-tagged proteins are visualized in green (middle panels) by anti-FLAG immunostaining and fluorescent secondary antibodies; merged images appear in yellow (right panels). Scale bars, 20 μm. Colocalization was quantified using Pearson’s correlation coefficient (R). Graphs represent mean ± SD from 20-30 cells per condition, based on ≥ 3 independent experiments.

To assess whether AGAP1 Influences the intracellular trafficking of HIV-1 Gag, we examined the localization of Gag-RFP in cells co-expressing AGAP1 constructs. In HEK 293T cells, expression of WT FLAG-AGAP1 induced a pronounced redistribution of Gag-RFP from the plasma membrane to intracellular compartments, often concentrated in perinuclear regions. In contrast, co-expression of the FLAG-AGAP1/R599K mutant did not significantly alter Gag-RFP localization, which remained primarily at the plasma membrane (Fig. 3C). Quantitative analysis revealed no significant differences in the degree of colocalization between Gag-RFP and either WT or mutant AGAP1 (Pearson’s R: 0.40 ± 0.14 for WT, 0.49 ± 0.10 for R599K). Similar results were observed in HeLa cells, where WT AGAP1 disrupted Gag-RFP targeting to the plasma membrane, while the AGAP1/R599K mutant had minimal effect (Fig. 3D). As before, colocalization between Gag-RFP and AGAP1 constructs did not differ significantly (Pearson’s R: 0.35 ± 0.11 for WT, 0.37 ± 0.07 for R599K). Parallel experiments using untagged Gag yielded a similar conclusion that expression of WT AGAP1, but not the R599K mutant, reduced Gag localization at the plasma membrane and induced its accumulation at internal compartment (Fig.S2).

These findings indicate that AGAP1 modulates Gag trafficking through its catalytic activity, rather than through direct interaction or stable colocalization with Gag. The inability of the GAP-deficient mutant to alter Gag localization underscores the requirement for spatially and temporally regulated Arf1 inactivation during the late stages of the HIV-1 replication cycle.

Together, these results demonstrate that AGAP1 negatively regulates HIV-1 particle release by modulating Gag trafficking in an Arf1 GAP activity-dependent manner.

### Arf6 modulates HIV-1 particle release and Gag subcellular localization

Arf6 is a small GTPase involved in endosomal membrane trafficking and actin cytoskeleton remodeling, and has been previously implicated in the late stages of the HIV-1 replication cycle through its regulation of Gag trafficking (25). To further define the role of Arf6 in HIV-1 particle assembly and release, HEK 293T cells were co-transfected with HIV-1 Gag-Pol and a constitutively active, GTP-locked mutant of Arf6 (HA-Arf6/Q67L) at increasing DNA ratios (1:1 and 1:2, relative to Gag-Pol), with empty vector (pcDNA3.1) added to equalize total DNA input. Immunoblot analysis of cell lysates and pelleted VLPs revealed a dose-dependent reduction in extracellular VLP production upon HA-Arf6/Q67L expression. Notably, intracellular levels of Pr55Gag and its processing intermediates were unaffected, indicating that Arf6 activation impairs a post-translational step, likely involving Gag trafficking or particle release, rather than Gag expression or proteolytic processing. Quantification of VLP release efficiency confirmed a significant reduction in particle release in the presence of activated Arf6 (Fig. 4B). These results suggest that constitutive Arf6 activation disrupts a late stage in HIV-1 assembly or budding.

**Fig. 4.**
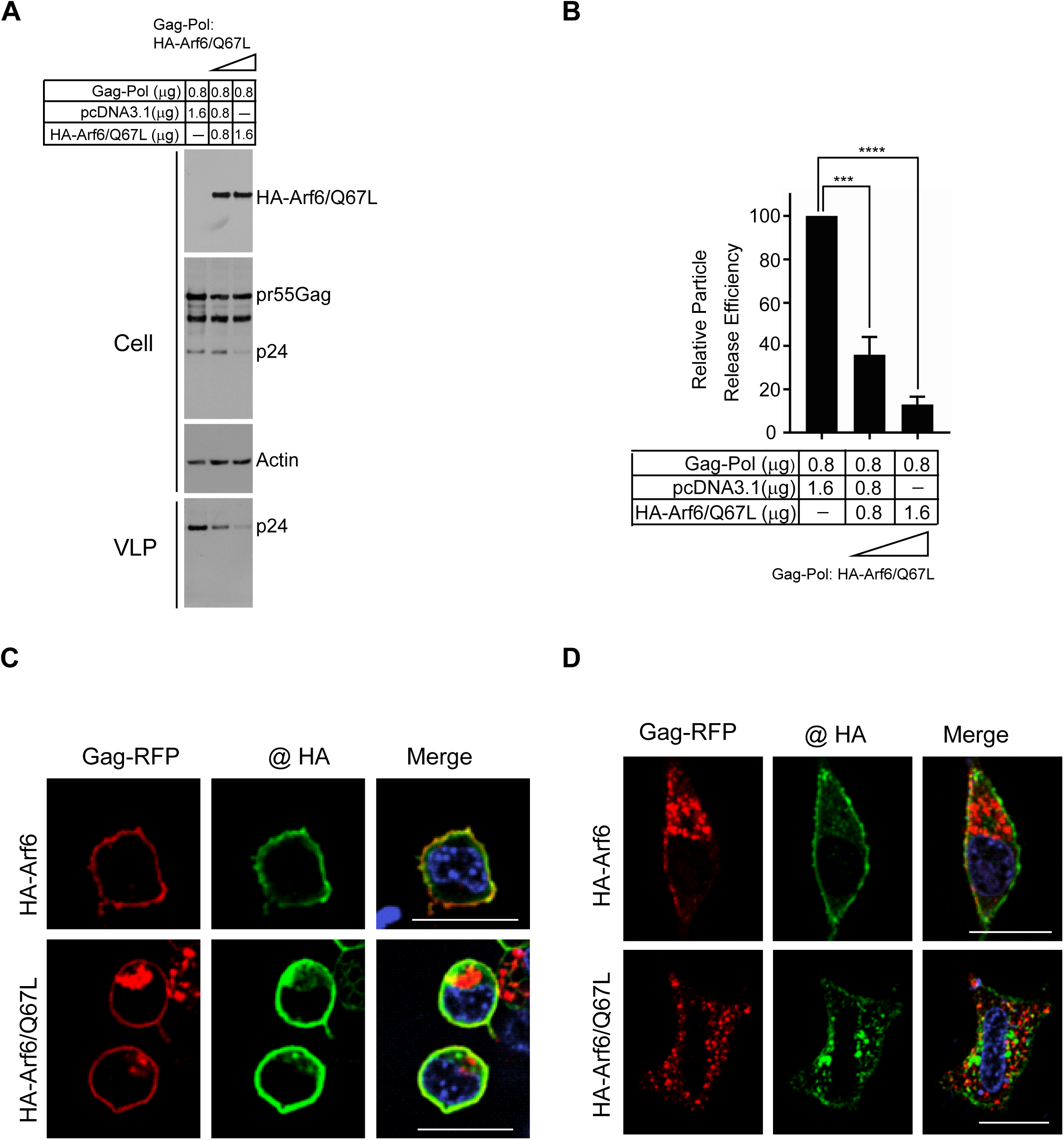
Arf6 modulates HIV-1 particle release. **A,** HEK 293T cells were co-transfected with HIV-1 Gag-Pol alone (lane 1) or with HA-Arf6/Q67L at increasing DNA ratios (lanes 2 and 3; 1:1 and 1:2, Gag-Pol: Arf6/Q67L). Total DNA input was equalized using empty vector (pcDNA3.1). Cell lysates were analyzed by immunoblotting with anti-HA, HIV-IgG and anti-actin antibodies. Pelleted VLPs were immunoblotted with HIV-IgG. **B,** Quantification of particle release efficiency from (A), normalized to control (lane 1). Data represent mean ± SD (n=3). ***, *p* < 0.001; ****, *p* < 0.0001. **C-D,** Confocal fluorescence microscopy of HEK 293T (C) and HeLa (D) cells co-expressing Gag-RFP (red) and either WT HA-Arf6 (top row) or HA-Arf6/Q67L (bottom row). HA-tagged proteins were detected by immunostaining with anti-HA antibodies and Alexa Fluor-conjugated secondary antibodies (green). Merged images show co-localization in yellow. Scale bars, 20 μm,

To determine the impact of Arf6 on Gag subcellular localization, we performed confocal fluorescence microscopy in HEK 293T co-expressing Gag-RFP and either WT HA-Arf6 or HA- Arf6/Q67L. At 24 h post-transfection, WT HA-ARf6 strongly colocalized strongly Gag-RFP at the plasma membrane, without overt changes in Gag distribution (Fig. 4C). In contrast, HA- Arf6/Q67L expression led to a marked redistribution of Gag-RFP to intracellular puncta, consistent with retention in endosomal or recycling compartments. Similar results were observed in HeLa cells, where WT HA-Arf6 co-expression supported Gag-RFP localization at both the plasma membrane and intracellular compartments. However, expression of HA-Arf6/Q67L resulted in predominant intracellular localization of Gag-RFP and reduced plasma membrane association (Fig. 4D). Similarly, Arf6/Q67L expression impaired the trafficking of untagged Gag to the plasma membrane (Fig. S3). Collectively, these data indicate that constitutively active Arf6 perturbs Gag trafficking and membrane targeting, thereby interfering with efficient HIV-1 particle assembly and release.

### Arf1-mediated trafficking pathways do not alter the subcellular localization or surface expression of BST-2

The antiviral function of BST-2 is tightly linked to its subcellular localization. Under steady-state conditions, BST-2 is distributed between the plasma membrane and intracellular compartments, including the TGN and recycling endosomes. Given the central role of Arf1 in mediating vesicle transport between Golgi and endosomes, we investigated whether modulation of Arf1 impacts the subcellular distribution of endogenous BST-2. HeLa cells were transfected with HA-tagged constructs encoding WT Arf1, Arf1/Q71L, or Arf1/T31TN. Cells were subsequently analyzed by immunofluorescence microscopy using antibodies against the HA epitope and endogenous BST-2. In all conditions, BST-2 exhibited its characteristic punctate distribution, with enrichment at the plasma membrane and a perinuclear compartment consistent with the TGN. Each Arf1 construct showed substantial colocalization with BST-2-positive compartments (Fig. 5A).

**Fig. 5.**
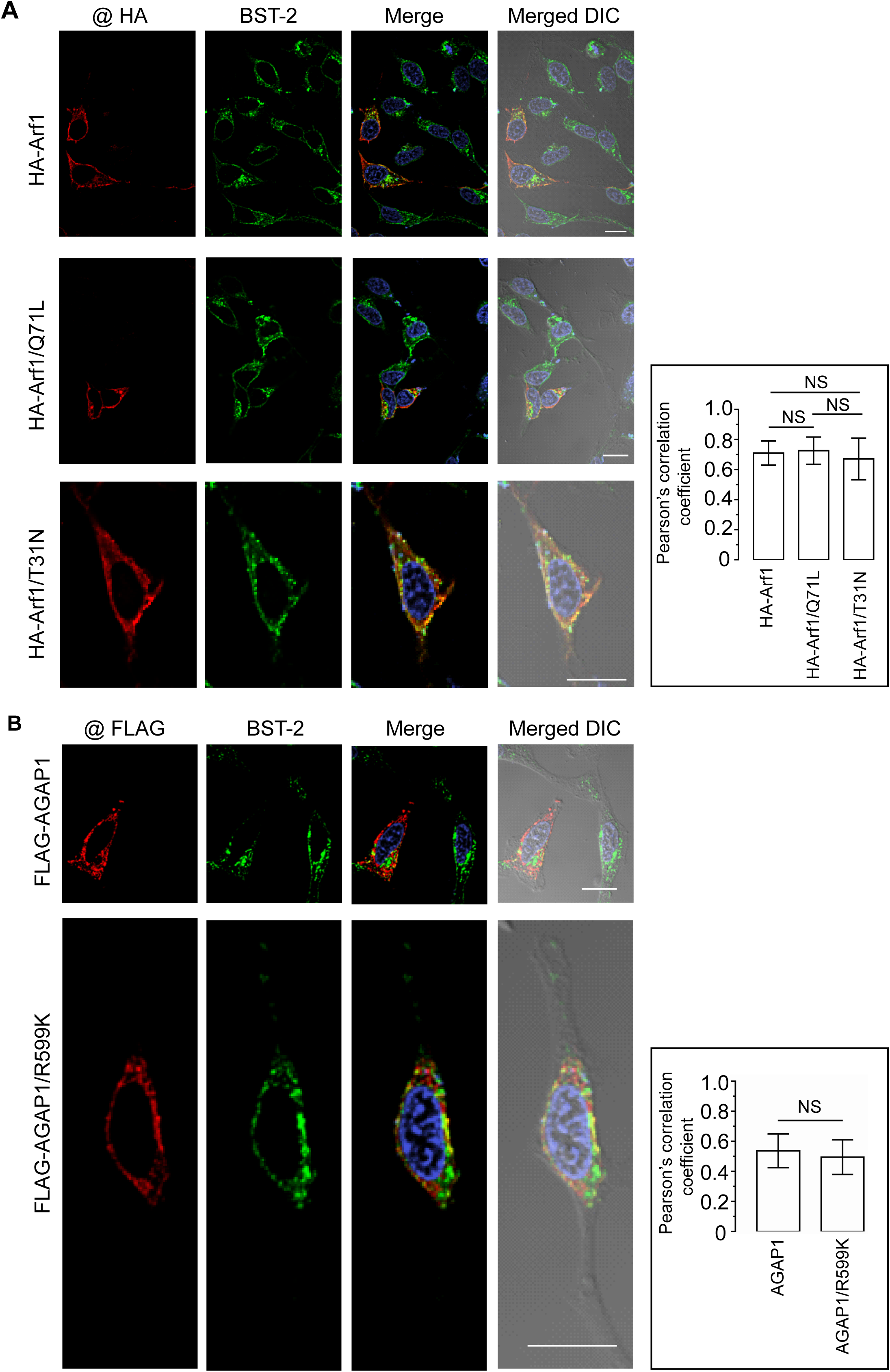
Disruption of Arf1 function does not alter BST-2 subcellular distribution. **A,** HeLa cells were transfected with HA-Arf1 (top row), HA-Arf1/Q71L (middle row), or HA-Arf1/T31N (bottom row). At 30 h post-transfection, cells were fixed, permeabilized, and stained with anti-HA and anti-BST-2 antibodies. HA-tagged proteins are shown in red (far-left panels), endogenous BST-2 in green (left panels), and the co-localized pixels in yellow (right panels). Differential interference contrast (DIC) images merged with confocal images are shown in the far-right panels. The graph on the right shows Pearson’s correlation coefficients quantifying co-localization between HA-tagged proteins and endogenous BST-2. Data represent mean ± SD from 20-25 cells, based on at least three independent experiments. Scale bars, 20 μm. **B,** HeLa cells were transfected with FLAG-AGAP1 (top row) or FLAG-AGAP1/R599K (bottom row), followed by immunostaining with anti-FLAG and anti-BST-2 antibodies. FLAG-tagged proteins are shown in red (far-left panels), BST-2 in green (middle panels), and the colocalization in yellow (right panels). DIC images merged with confocal images are shown in the far-right panels. Pearson’s correlation coefficients for colocalization between FLAG-tagged proteins and BST-2 are shown in the accompanying graph (right). Data represent mean ± SD from 20-25 cells, based on at least three independent experiments. Scale bars, 20 μm.

Quantification of Pearson’s correlation coefficients from > 30 cells per condition revealed high colocalization values: 0.71 ± 0.08 for WT Arf1, 0.73 ± 0.09 for Arf1/Q71L, and 0.67 ± 0.14 for Arf1/T31N. No statistically significant differences were observed, indicating that Arf1 activity does not alter the subcellular localization of BST-2. We next assessed whether AGAP1 modulates BST-2 distribution. HeLa cells were transfected with FLAG-tagged wild-type AGAP1 or a mutant (R599K) and stained for FLAG and endogenous BST-2. Both constructs exhibited a perinuclear distribution and partial colocalization with BST-2 (Fig. 5B). However, BST-2 maintained its distribution between the plasma membrane and TGN regardless of AGAP1 expression. Pearson’s correlation coefficients were 0.54 ± 0.11 for WT AGAP1 and 0.50 ± 0.11 for the R599K mutant, with no significant difference between groups, suggesting that AGAP1 does not significantly alter BST-2 localization.

To determine whether Arf1 affects total or surface BST-2 expression levels, we performed immunoblotting of total cell lysates and flow cytometric analysis of surface BST-2 in transfected HeLa cells. Western blot analysis revealed no changes in total BST-2 protein levels upon expression of Arf1 WT, Q71L, or T31N, as normalized to actin (Fig. 6A). Flow cytometry of GFP-positive cells co-transfected with Arf1 constructs showed that surface BST-2 expression remained comparable to empty vector controls (Fig. 6B).

**Fig. 6.**
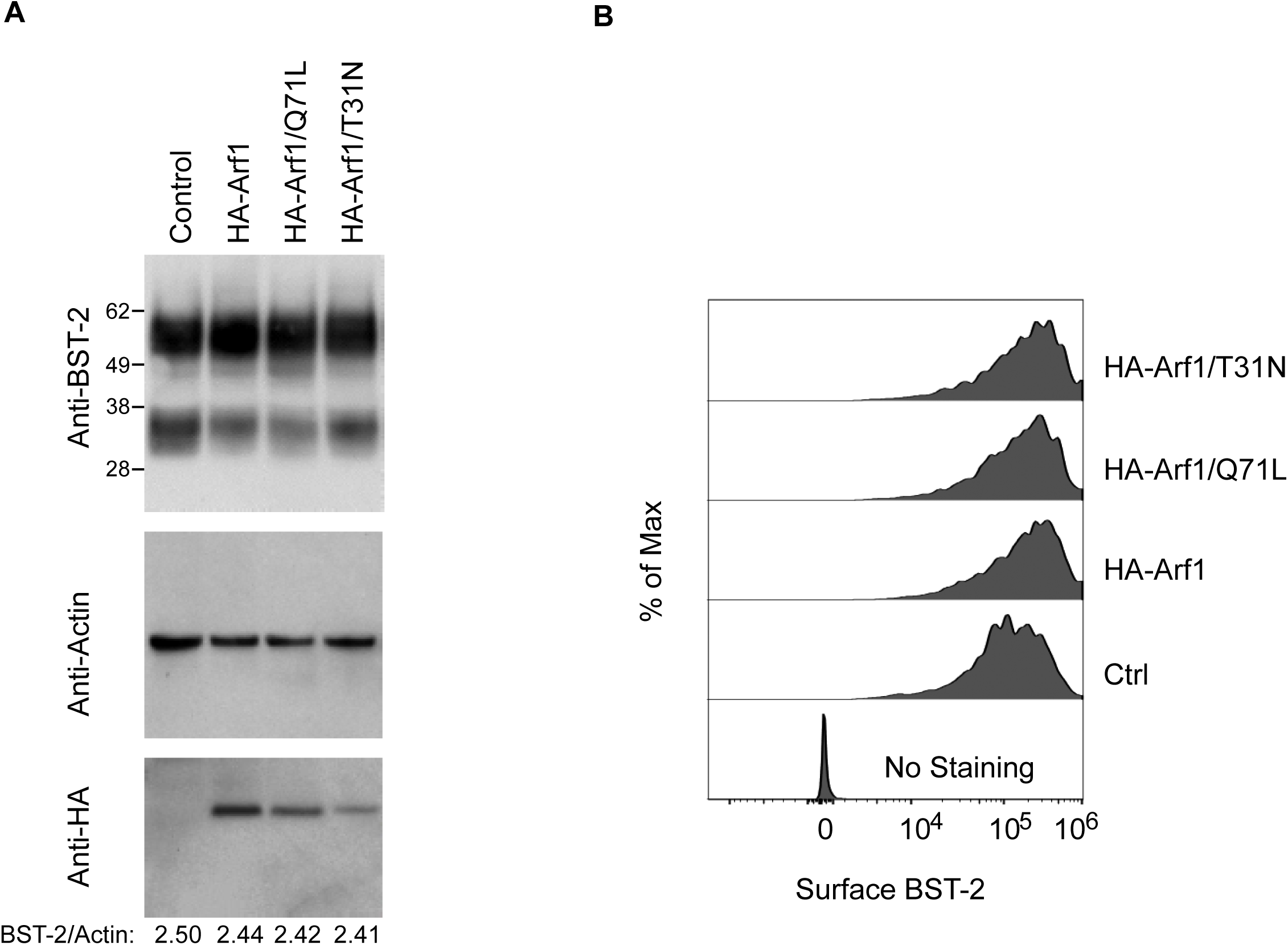
Impairment of Arf1 function does not change BST-2 expression. **A,** HeLa cells were transfected with pcDNA3.1 (lane 1), HA-Arf1 (lane 2), HA-Arf1/Q71L (lane 3), or HA-Arf1/T31N (lane 4). Cell lysates were analyzed by immunoblotting with anti-HA, anti-BST-2, and anti-actin antibodies. Densitometric quantification of BST-2 levels normalized to actin is shown below. Data shown are representative of three independent experiments. **B,** HeLa cells were co-transfected with GFP and either pcDNA3.1, HA-Arf1, HA-Arf1/Q71L, or HA-Arf1/T31N. Surface expression of BST-2 on GFP-positive cells was assessed by flow cytometry.

Collectively, these data indicate that neither Arf1 activity state nor modulation of Arf1 by AGAP1 affects BST-2 subcellular localization, surface expression, or overall abundance. These findings suggest that the Arf1/AGAP1 regulatory axis does not modulate the antiviral activity of BST-2 through changes in its intracellular distribution or surface availability.

### Arf6 does not modulate BST-2 localization or expression

To determine whether Arf6 influences BST-2 localization or expression, we transfected HeLa cells with HA-tagged WT Arf6 or Arf6/Q67L, and performed confocal immunofluorescence microscopy 30 h post-transfection. Cells were stained with anti-HA and anti-BST-2 antibodies to visualize Arf6 expression and endogenous BST-2 distribution. Both HA-Arf6 and HA-Arf6/Q67L exhibited cytoplasmic localization with enrichment at the plasma membrane and perinuclear compartments, partially overlapping with BST-2. However, neither construct caused a detectable redistribution of BST-2 from its typical localization in the TGN, recycling endosomes, and cell surface (Fig. 7A). Quantitative colocalization analysis was performed to assess the degree of overlap between Arf6 and BST-2. WT HA-Arf6 showed moderate colocalization with BST-2 (R=0.46 ± 0.14), while HA-Arf6/Q67L exhibited slightly higher colocalization (R=0.59 ± 0.05), based on analysis of 30-35 cells across three independent experiments. These values indicate moderate spatial association but not functional co-trafficking or sequestration of BST-2.

**Fig. 7.**
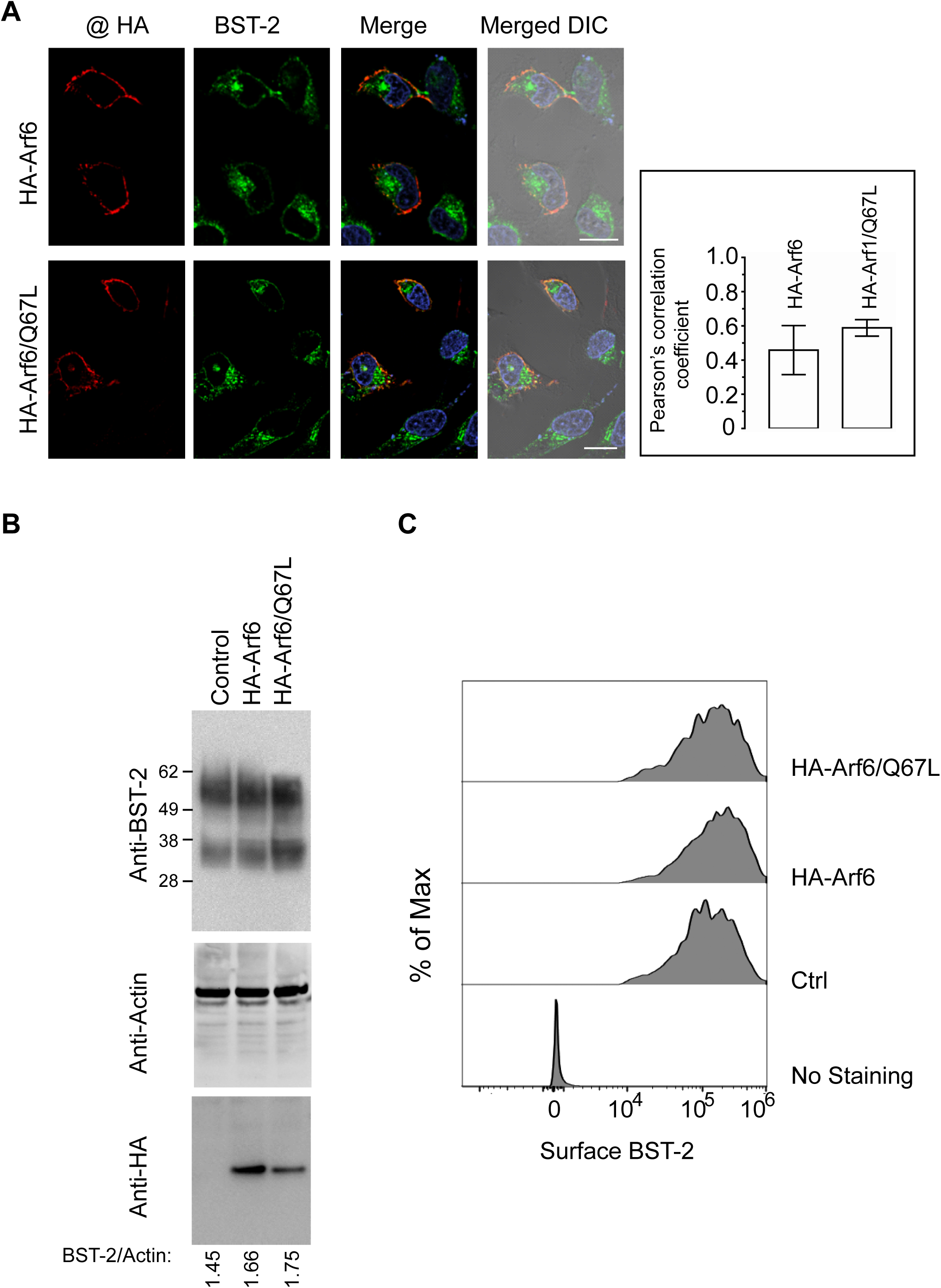
Arf6 function does not affect BST-2 expression or localization. **A,** HeLa cells were transfected with HA-Arf6 (top row) or HA-Arf6/Q67L (bottom row), fixed, permeabilized, and stained with anti-HA and anti-BST-2 antibodies. Confocal images show HA-tagged proteins in red (far-left panels), BST-2 in green (left panels), and co-localized pixels in yellow (right panels). DIC images merged with fluorescence channels are shown in the far-right panels. The graph on the right shows Pearson’s correlation coefficients quantifying colocalization between HA-tagged proteins and BST-2. Data represent mean ± SD from 20-25 cells, based on three independent experiments. **B,** HeLa cells were transfected with pcDNA3.1 (lane 1), HA-Arf6 (lane 2), or HA-Arf6/Q67L (lane 3). Cell lysates were analyzed by immunostaining with anti-BST-2, anti-actin, and anti-HA antibodies. Densitometric ratios of BST-2 signal intensities normalized to corresponding actin levels are shown below. Data shown are representative from three independent experiments. **C,** HeLa cells were co-transfected with GFP and either pcDNA3.1, HA-Arf6, or HA-Arf6/Q67L. Surface expression of BST-2 on GFP-positive cells was analyzed by flow cytometry.

To assess whether Arf6 expression affects the overall abundance of BST-2, we performed immunoblot analysis of whole-cell lysates from cells transfected with either vector control, HA-Arf6, or HA-Arf6/Q67L. Immunoblotting with anti-BST-2 and anti-actin antibodies revealed no significant difference in BST-2 levels between groups (Fig. 7B). Densitometric quantification of BST-2 signal intensity normalized to actin confirmed the absence of significant changes.

Additionally, we assess the surface expression of BST-2 by flow cytometry. HeLa cells were co-transfected with GFP and either vector control, HA-Arf6, or HA-Arf6/Q67L, and surface BST-2 levels were measured in the GFP-positive population using an anti-BST-2 antibody that recognizes the extracellular domain. Flow cytometry analysis showed that surface BST-2 expression remained unchanged in the presence of either wild-type or mutant Arf6 compared to control (Fig. 7C), further indicating that Arf6 activity does not alter BST-2 trafficking to the plasma membrane. Taken together, these data demonstrate that Arf6, like Arf1, does not significantly influence the subcellular distribution or expression level of BST-2, supporting the conclusion that BST-2 is not regulated by Arf6-mediated membrane trafficking pathways.

## DISCUSSION

This study identifies a critical role for the small GTPase Arf1 and its regulatory machinery in modulating the late stages of the HIV-1 replication cycle. Specifically, we demonstrate that dynamic cycling between the GTP- and GDP-bound states of Arf1 is required for efficient trafficking of the HIV-1 Gag polyprotein to the plasma membrane and for productive virion release. Disruption of this cycle, either through constitutive activation (Q71L) or inactivation (T31N), impairs Gag membrane association and significantly reduces VLP output. These findings underscore the necessity for tightly coordinated Arf1 activity to support viral particle assembly and reinforce the broader principle that small GTPases act as molecular switches that govern vesicle-mediated trafficking.

Arf1 is a well-characterized regulator of intracellular membrane trafficking, particularly at the Golgi apparatus and early endosomal compartments, where it recruits coat proteins (e.g., COPI), lipid-modifying enzymes, and contributes to membrane curvature (39–43). While its cellular functions are well understood, relatively few studies have linked Arf1 to retroviral replication (24,44). Our data now provide direct evidence that Arf1 activity modulates the subcellular localization of Gag, a critical determinant of HIV-1 assembly. Both membrane flotation assays (Fig. 2) and immunofluorescence imaging (Fig.1, Fig.S1) demonstrate that disrupting Arf1 cycling impairs Gag membrane association and prevents its accumulation at the plasma membrane, thereby compromising VLP release. Interestingly, Arf1 itself shows limited colocalization with Gag, suggesting an indirect regulatory mechanism (Fig. 1). We propose that Arf1 shapes the trafficking environment, potentially by influencing membrane identity or lipid composition at the TGN or recycling endosomes, to promote efficient Gag sorting and delivery. Alternatively, Arf1 may regulate the generation of specific phosphoinositide lipids, such as PI(4)P or PI(4,5)P2, which are critical for Gag membrane binding (45–47). Indeed, Arf1 is known to recruit lipid kinases and phosphatases that help establish organelle-specific lipid profiles (48,49), and Gag has been shown to preferentially associate with PI(4,5)P2 at the plasma membrane (25).

A major insight from this study is the identification of AGAP1, an Arf1-specific GTPase-activating protein, as a negative regulator of HIV-1 assembly. AGAP1 promotes GTP hydrolysis on Arf1, thereby converting it to inactive GDP-bound form (37). Overexpression of AGAP1 markedly reduced VLP release and caused Gag mislocalization, phenocopying the effects of the GDP-locked Arf1/T31N mutant. In contrast, a GAP-deficient AGAP1 mutant (R599K) failed to affect Gag trafficking or virion production, confirming that the catalytic activity of AGAP1 is essential for its regulatory function (Fig.3, Fig. S2). These data support a model in which the precise spatial and temporal control of Arf1 inactivation, likely coordinated by AGAP1 at specific intracellular trafficking hubs, is required for efficient Gag transport to the plasma membrane. Excessive or premature inactivation of Arf1, as seen with AGAP1 overexpression, may redirect Gag-containing vesicles into nonproductive pathways or disrupt vesicle formation altogether.

Furthermore, our results suggest that GGAs and AGAP1 act in a coordinated sequence within the Arf1-mediated trafficking pathway, particularly during clathrin-coated vesicle formation and cargo transport. GGAs serve as adaptor proteins that link activated Arf1 to clathrin, promoting vesicle budding. Arf1 likely dissociates from GGAs prior to its inactivation by AGAP1, supporting a sequential model in which GGAs and AGAP1 compete for overlapping binding sites on Arf1 but act at different stages (50–52). Consistent with this, previous studies have showed that GGA overexpression disrupts intracellular sorting and impairs retroviral particle release (24,44).

Notably, both persistent activation and over-inactivation of Arf1 inhibit viral assembly, revealing a “Goldilocks” principle of GTPase function, where balanced cycling between active and inactive states is critical. Similar paradigms have been described for Rab and Rho GTPases, whose cycles of nucleotide exchange and hydrolysis are tightly regulated to ensure vesicle formation, transport, docking, and fusion(53–58). Our findings now extend this model to Arf1 in the context of HIV-1, highlighting an unexpected vulnerability of the viral assembly process to disruptions in host membrane trafficking networks.

We further demonstrate that Arf6, a related GTPase associated with endosomal membranes and the plasma membrane (59,60), also disrupts HIV-1 Gag trafficking when constitutively activated. Expression of the GTP-locked Arf6/Q67L mutant caused mislocalization of Gag from the plasma membrane to intracellular compartments and suppressed VLP production (Fig.4, Fig. S3). These observations are consistent with prior studies implicating Arf6 in actin remodeling, endosomal recycling, and lipid composition regulation (61–63)−processes that converge on the late stages of viral budding. It is plausible that sustained Arf6 activation disrupts actin-dependent transport, alters lipid domains, or interferes with vesicle scission at the plasma membrane. In contrast, WT Arf6 did not affect Gag localization (Fig. 4), indicating that the observed defects result from aberrant GTPase activity rather than simple overexpression.

Importantly, our data rule out the possibility that Arf1 or Arf6 regulate HIV-1 release via modulation of BST-2, a host restriction factor that inhibits virion detachment by physically tethering nascent virions at the cell surface (30,33,64). Although both Arf GTPases showed moderate colocalization with BST-2-positive compartments, neither altered the expression level, surface abundance, nor steady-state distribution of endogenous BST-2 (Fig.5, 6, 7). AGAP1 similarly failed to impact BST-2 localization (Fig.5). These findings suggest that Arf1 and Arf6 do not modulate BST-2 function through trafficking changes and that the observed defects in VLP release are BST-2-independent. This conclusion is consistent with previous studies indicating that HIV-1 Vpu is the principal antagonist of BST-2, promoting its degradation and downregulation (34,65–69).

Taken together, our results uncover a previously unappreciated requirement for Arf1 cycling in the late stages of the HIV-1 life cycle. Arf1 activity governs Gag trafficking, membrane association, and particle release, and this process is exquisitely sensitive to deviations in GTPase activity. AGAP1 serves as a key negative regulator by modulating Arf1 inactivation, and constitutive Arf6 signaling similarly disrupts Gag trafficking, underscoring the broader relevance of Arf family GTPases in retroviral assembly. Future studies will be necessary to identify the downstream effectors of Arf1 and Arf6 involved in this pathway, to map the relevant membrane compartments supporting Gag trafficking, and to determine whether pharmacological targeting of Arf-regulatory circuits could provide new avenues to disrupt HIV-1 replication.

## EXPERIMENTAL PROCEDURES

### Cell Cultures and Transfections

HeLa (ATCC CCL2) and HEK 293T (ATCC CRL-3216) cells were obtained from the American Type Culture Collection (ATCC) and maintained in Dulbecco’s Modified Eagle’s Medium (DEME; high glucose; Thermo Fisher Scientific) supplemented with 2 mM L-glutamine, 10% fetal bovine serum (FBS; Gibco), 100 U/ml penicillin, and 100 μg/ml streptomycin at 37°C in a humidified incubator with 5% CO2. HeLa cells were transfected using either Lipofectamine 2000 (Thermo Fisher Scientific) or TransIT-HeLaMONSTER (Mirus Bio) according to the manufacturers’ instructions. HEK 293T cells were transfected using either polyethyleneimine (PEI, Sigma-Aldrich), calcium phosphate precipitation, or X-tremeGENE HP DNA transfection reagent (Sigma-Aldrich).

### Plasmids

An expression plasmid encoding HA-tagged full-length Arf1 was kindly provided by Juan Bonifacino. The following plasmids were obtained from Thomas Roberts via Addgene, including pcDNA3-HA-Arf1/Q71L (plasmid #10800) encoding HA-tagged Arf1 mutation with Q71L substitution, pcDNA3-HA-Arf1/T31N (plasmid #10801) encoding HA-tagged Arf1 mutation with T31N substitution, pcDNA3-HA-Arf6 (plasmid # 10802) encoding HA-tagged full-length Arf6, and pcDNA3-HA-Arf6/Q67L (plasmid # 10803) encoding HA-tagged Arf6 mutation with Q67L substitution (70). The HIV-1 Gag-Pol expression plasmid pGPCINS was provided by Xiao-Fang Yu (71), and the Gag-RFP construct was a kind gift from Akira Ono (72). FLAG-tagged AGAP1 (WT and R599K mutant) constructs were obtained from Paul Randazzo (73).

### Flow Cytometry

HeLa cells were co-transfected with GFP and either HA-tagged Arf1, Arf1/Q71L, Arf1/T31N, Arf6, or Arf6/Q67L constructs. At 48 h post-transfection, cells were stained with APC-conjugated anti-human BST-2 antibodies (BioLegend) and analyzed using a Luminex Amnis CellStream flow cytometer. Data were processed and analyzed with FlowJo software (BD Biosciences).

### Immunofluorescence Microscopy

Confocal immunofluorescence microscopy was performed as described previously (74). For single-staining experiments, HA- or FLAG-tagged proteins were detected using mouse anti-HA (Abcam) or anti-FLAG (Sigma-Aldrich) antibodies, followed by Alexa Fluor 488-conjugated goat anti-mouse secondary antibodies (Thermo Fisher Scientific). For double-staining, Alexa Fluor 546-conjugated goat anti-mouse secondary antibodies (Thermo Fisher Scientific) were used to visualize HA- or FLAG-tagged proteins. Endogenous BST-2 was detected using rabbit anti-BST-2 antibodies (Abcam) and Alexa Fluor 488-conjugated goat anti-rabbit secondary antibodies (Thermo Fisher Scientific). HIV-1 Gag was detected using rabbit anti-p17 antibodies (NIH AIDS Reagent Program) and Alexa Fluor 488-conjugated goat anti-rabbit secondary antibodies. Images were acquired on a Nikon AIR confocal microscope and analyzed using NIS-Elements AR software (Nikon Instruments Inc.).

### VLP Production and Purification

HEK 293T Cells were co-transfected with HIV-1 Gag-Pol and either empty vector (pcDNA3.1), HA-Arf1, HA-Arf1/Q71L, HA-Arf1/T31N, FLAG-AGAP1, or HA-Arf6/Q67L. At 48 h post-transfection, supernatants were collected, cleared by centrifugation at 500 × *g* for 10 min, filtered through a 0.45µm membrane, and centrifuged through a 20% sucrose cushion at 28, 000 × *g* for 2 h at 4°C. Pelleted VLPs were resuspended and analyzed by western blot.

### Membrane Flotation Assay

A modified membrane flotation assay was used to assess HIV-1 Gag membrane association (75). HEK 293T cells were co-transfected with HIV-1 Gag and either pcDNA3.1, HA-Arf1, HA-Arf1/Q71L, or HA-Arf1/T31N. At 48 h post-transfection, cells were harvested using trypsin-EDTA, resuspended in hypotonic buffer (10 mM Tris-HCl, pH 8.0, supplemented with protease inhibitors), and incubated on ice for 15 min. Cells were lysed by Dounce homogenization and lysates were adjusted to 0.1 M NaCl. After clearing by centrifugation at 1,000 × *g* for 10 min at 4°C, the post-nuclear supernatant was adjusted to 50% (v/v) iodixanol from a 60% stock (Axis-Shield) and overlaid with 40%, 30%, and 10% iodixanol solutions prepared in buffer (0.85% NaCl, 10 mM Tricine-NaOH, pH7.4). Gradients were centrifuged at 35,000 rpm for 16 h at 4°C in an SW41Ti rotor (Beckman Coulter). Fractions were collected from the top and analyzed by western blot using anti-p24 (BEI Resources) and anti-HA (Abcam) antibodies.

### Statistical Analysis

Quantitative data represent the mean ± standard deviation (SD) from at least three independent experiments unless otherwise noted. Statistical significance was determined using unpaired two-tailed Student’s *t*-tests in GraphPad Prism v10 (GraphPad Software). A *p*-value < 0.05 was considered statistically significant.

## Data Availability

All data supporting the findings of this study are included within the article and its supplementary materials. Additional raw data and reagents are available from the corresponding author upon reasonable request.

## Acknowledgements

We thank Olga Korolkova and Qiujia Shao for their technical assistance with confocal microscopy and flow cytometry. The following reagent was obtained through BEI Resources, NIAID, NIH: monoclonal anti-HIV-1 p24 antibody (clone 183-H12-5C; Cat# ARP-3537), contributed by Drs. Bruce Chesebro and Kathy Wehrly. The following reagent was obtained through the NIH AIDS Research and Reference Reagent Program, Division of AIDS, NIAID, NIH: rabbit antiserum against HIV-1 p17 protein (Cat# ARP-4811), contributed by Drs. Paul Spearman and Lingmei Ding.

## Funding

This research was supported in part by the National Institutes of Health (NIH) grant R01 AI1557764 (to X.D.), the Tennessee Center for AIDS Research (CFAR) grant P30 AI110527, and the Research Centers in Minority Institutions (RCMI) grant U54 MD007586.

## Competing Interests

The authors declare that they have no conflicts of interest with the contents of this article. competing interests.

## Supporting Information

This article contains supporting information.

## Notes

### Competing Interest Statement

The authors have declared no competing interest.

