## Supplemental Figures 1-3 for "Arf GTPases Define BST-2-Independent Pathways for HIV-1 Assembly and Release"

Figure S1

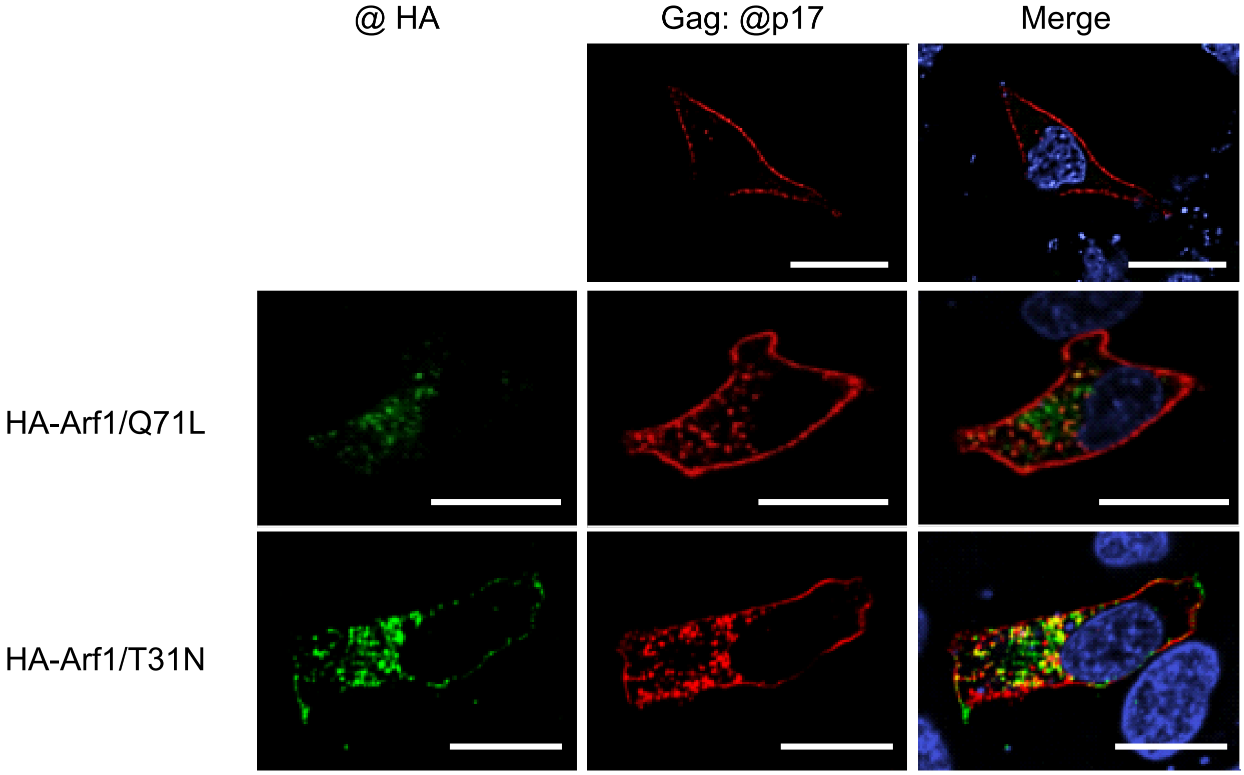


**Fig. S1. Arf1 regulates HIV-1 Gag subcellular localization.** HeLa cells were co-transfected with HIV-1 Gag alone (top row), Gag with HA-Arf1/Q71L (middle row), or Gag with HA-Arf1/T31N (bottom row). At 24 h post-transfection, cells were fixed, permeabilized, and stained with anti-p17 (to detect Gag) and anti-HA antibodies. HA-tagged proteins are shown in green (left panels), Gag in red (middle panels), and merged images in yellow (right panels). Scale bars, 20 um.

Figure S2

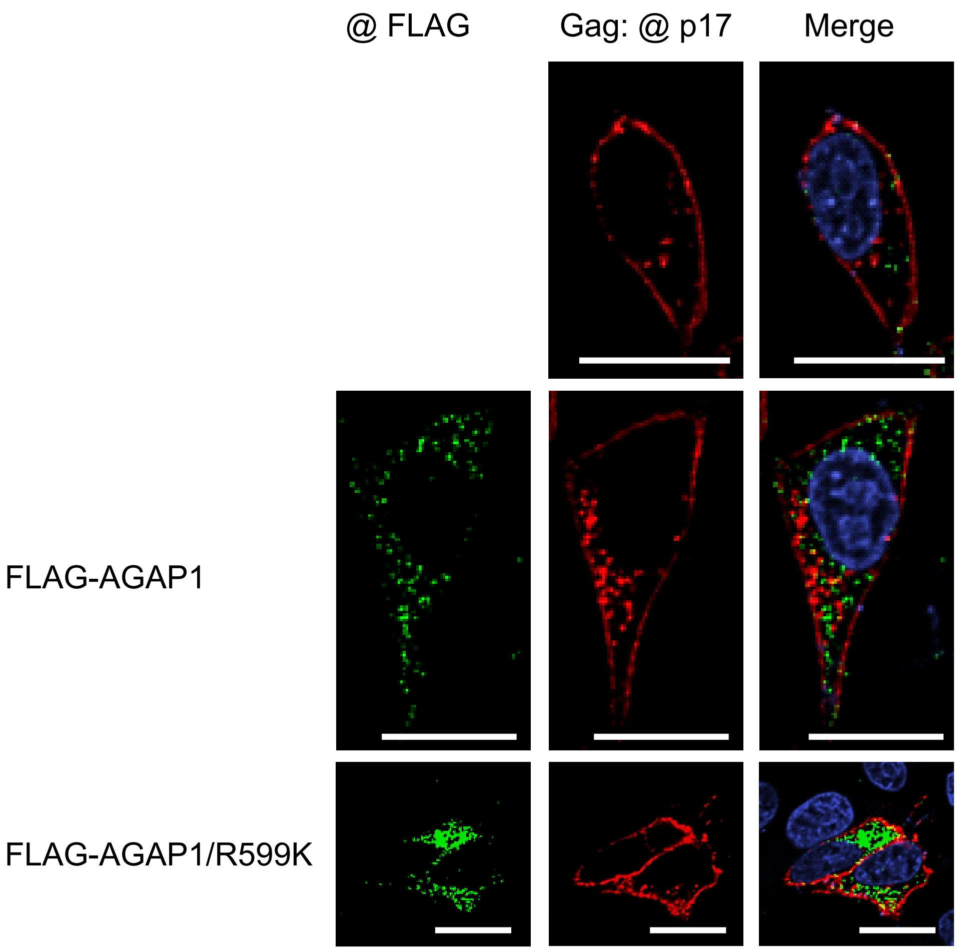


**Fig. S2. AGAP1 modulates HIV-1 Gag subcellular localization.** HeLa cells were co-transfected with HIV-1 Gag alone (top row), Gag with FLAG-AGAP1 (middle row), or Gag with FLAG-AGAP1/R599K (bottom row). At 24 h post-transfection, cells were fixed, permeabilized, and stained with anti-p17 (to detect Gag) and anti-FLAG antibodies. FLAG-tagged proteins are shown in green (left panels), Gag in red (middle panels), and merged images in yellow (right panels). Scale bars, 20 um.

Figure S3

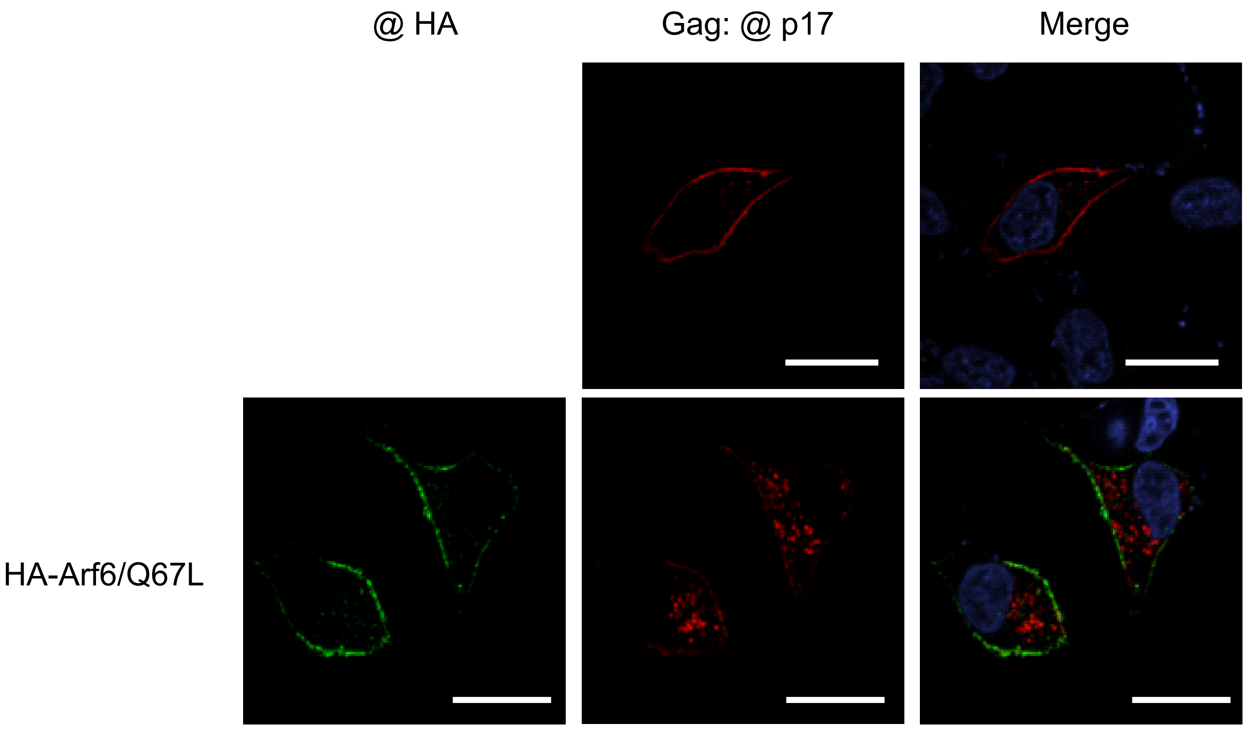


**Fig. S3. Arf6 regulates the subcellular localization of HIV-1 Gag.** HeLa cells were co-transfected with HIV-1 Gag alone (top row),  or Gag with HA-Arf6/Q67L(bottom row). At 24 h post-transfection, cells were fixed, permeabilized, and stained with anti-p17 (to detect Gag) and anti-HA antibodies. HA-Arf6/Q67L is shown in green (left), Gag in red (middle panels), and merged images in yellow (right panels). Scale bars, 20 um.
